# Functional characterization of a neuropeptide receptor exogenously expressed in *Aplysia* neurons

**DOI:** 10.1101/2022.02.14.480444

**Authors:** Guo Zhang, Shi-Qi Guo, Si-Yuan Yin, Wang-Ding Yuan, Ping Chen, Ji-Il Kim, Hui-Ying Wang, Hai-Bo Zhou, Abraham J. Susswein, Bong-Kiun Kaang, Jian Jing

**Affiliations:** State Key Laboratory of Pharmaceutical Biotechnology, Institute for Brain Sciences, School of Life Sciences, Nanjing University, Nanjing, Jiangsu 210023, China; School of Biological Sciences, Seoul National University, 1, Gwanak-ro, Gwanak-gu, Seoul, 08826, Korea; School of Electronic Science and Engineering, Nanjing University, Nanjing, Jiangsu 210023, China; Peng Cheng Laboratory, Shenzhen 518000, China; The Mina and Everard Goodman Faculty of Life Sciences, Bar Ilan University, Ramat Gan 52900, Israel; Department of Neuroscience, Icahn School of Medicine at Mount Sinai, New York, NY 10029, USA

## Abstract

Neuropeptides act mostly on a class of G-protein coupled receptors, and play a fundamental role in the functions of neural circuits underlying behaviors. However, the functions of neuropeptide receptors are poorly understood. Here, we used the mollusc model system *Aplysia* and microinjected the exogenous neuropeptide receptor apATRPR (*Aplysia* Allatotropin-related peptide receptor) with an expression vector (pNEX3) into *Aplysia* neurons that did not express the receptor. Physiological experiments demonstrated that apATRPR could mediate the excitability increase activated by its ligand, apATRP (*Aplysia* Allatotropin-related peptide), in the *Aplysia* neurons that now express the receptor. This study provides the first definitive evidence for a physiological function of a neuropeptide receptor in molluscan animals.

## Introduction

Neuropeptides are the most diverse class of neurotransmitters/neuromodulators, which mostly act on G-protein coupled receptors (GPCRs). The diversity arises in part from the possibility that a single neuropeptide precursor can generate multiple forms of active peptides, and a peptide can act on multiple GPCRs, which in turn might function through different signaling pathways [1, 2]. Consequently, it has been challenging to study peptide signaling systems in vertebrates. Thus, relatively simple model systems, such as *Aplysia* [3-28], with identifiable neurons in well-defined neural circuits [3-10, 12, 14-17, 21, 22, 24-36] have been used in these studies. Earlier studies in model systems have focused on identifying neuropeptides and their bioactivity [2, 34, 35, 37-47]. Recently, growing genetic information in model systems is becoming available and has facilitated studying of both neuropeptides and their receptors [36, 48-52]. In the latter studies [49, 50], a common approach is to express putative GPCRs in a cell line, and test activity of potential ligands on the receptor. Then the receptor expression in the CNS and physiological and/or circuit activity of the ligands are demonstrated. If the receptor activity of the ligands in the cell line matches their physiological activity in the CNS, it is used as evidence that the receptor functions in the CNS. However, given that a peptide might act on multiple receptors, it is necessary to demonstrate that the identified GPCR actually displays the proper physiological activity in native neurons. In this paper, we have used an expression vector [3-5], to develop a method that expresses a peptide GPCR in native *Aplysia* neurons, and examine whether the GPCR could show a physiological activity. Our research utilizes *Aplysia* allatotropin (apATRP) [39] and its receptor apATRPR [50] as an example.

The neuropeptide allatotropin was first found in tissues of corpora allata in the insect *Manduca Sexta*, and it stimulated the secretion of juvenile hormone [53]. Subsequently, Allatotropin-related peptides have been characterized in animals across phyla, including Arthropoda [54], Annelida [48] and Mollusca [39] with multiple functions in different behaviors, including feeding. The allatotropin receptor was originally characterized in *Bombyx mori* [55], followed by more insect allatotropin receptors, such as in *Aedes aegypti* [56] and *Tribolium castaneum* [57]. Additionally, two allatotropin receptors in the annelid *Platynereis* [48] and one in *Aplysia* [50] were characterized. Interestingly, although allatotropin in the protostomes and orexin/hypocretin in vertebrates/deuterostomes [58] display no obvious similarity other than the amidated C-terminal, recent work has shown that their receptors are orthologous to each other based on phylogenetic analyses, supporting the evolutionary homology of allatotropin and orexin signaling systems [59, 60].

In *Aplysia*, apATRP (GFRLNSASRVAHGY-NH2) functions in the feeding network have been extensively characterized. apATRP targets B61/B62 motor neurons in the buccal ganglia by enhancing their excitability [39]. B61/B62 is a site of plasticity after learning that food is inedible [26]. Recently, several ligands, including apATRP, were found to activate apATRPR in CHO-K1 cells transiently transfected with apATRPR. Importantly, the pattern of activations of these ligands in the cell line matches their actions on B61/62 excitability, suggesting that apATRPR likely functions in the *Aplysia* CNS [50]. However, it is unknown whether apATRPR mediates the excitability increase in native *Aplysia* neurons. This is a critical question, given that there might be multiple apATRP receptors in *Aplysia*. Here, we sought to determine whether apATRP is sufficient to mediate the ligand effect on native neurons by evaluating the ability of apATRP to activate apATRPR in *Aplysia* neurons that do not originally express apATRPR.

To express apATRPR in neurons, we used a plasmid vector, pNEX (plasmid for neuronal expression), which is a reliable and effective method to express exogenous proteins in cultured *Aplysia* neurons [5, 61]. Initially, pNEX was constructed with the AK01a gene (shaker K^+^ channel), and microinjected into *Aplysia* neurons, which demonstrated that the AK01a channel could modulate the firing of the injected neuron and regulate synaptic interactions [61]. Subsequent studies used pNEX to overexpress the target proteins and elucidated molecular mechanisms of synaptic plasticity [62-68]. Notably, small molecule neurotransmitter receptors (1α metabotropic glutamate receptor [69]) and biogenic amine receptors (octopamine receptor) [64]), which are G-protein coupled receptors, were also expressed in *Aplysia* neurons through the vector pNEX. However, to date, no studies have applied this technique to neuropeptide GPCRs. Here, we used pNEX3 to successfully express apATRPR in *Aplysia* neurons that originally did not respond to ATRP. After expression, the neurons showed excitability increase in response to apATRP, indicating that apATRPR can mediate the excitability increase.

## Materials and methods

### Construction of plasmids

pNEX has two members, pNEXδ [3, 62, 63, 66, 70-74] or pNEX3 [5, 67-69, 75, 76]. There are eight more enhancers in pNEX3 than pNEXδ, making pNEX3 more effective in expressing a gene [5]. We chose to use pNEX3. Previously, plasmid vectors expressing the pNEX3 gene were constructed in *Aplysia* neurons [3, 5], and overexpressed proteins in a specific neuron [3, 77, 78]. The plasmid pNEX3-EGFP was derived from the earlier work [73], and the plasmid pcDNA3.1-apATRPR was a gift from Dr. Checco at the University of Illinois. To generate the expression plasmid pNEX3-apATRPR, the apATRPR gene was ligated with the vector pNEX3. First, the EGFP gene was digested from the BamHI-KpnI restriction fragment of pNEX3 and separated by agarose gel electrophoresis. Second, the apATRPR gene was added to the restriction sites of BamHI and KpnI at the 5’ and 3’ ends (forward primer: CGCGGATCCATGGGGTCGAACGATACATTC; reverse primer: GGGGTACCTCAGATGCTGGCGAGAGTGACCTC), respectively, by performing polymerase chain reaction (PCR) with the pcDNA3.1-apATRPR plasmid as the template. Then, the target gene, apATRPR, and the vector, pNEX3, were ligated using T4 DNA ligase. All plasmid DNAs used in microinjection were prepared by a standard maxi-prep procedure using an EndoFree Maxi Plasmid Kit.

### Electrophysiology

*Aplysia californica* (100−300 g) were purchased from Marinus Scientific (Long Beach, CA). *Aplysia* are hermaphroditic (i.e., each animal has functioning male and female reproductive organs). Animals were kept in an aquarium containing aerated and filtered artificial seawater (Instant Ocean, Aquarium Systems Inc., Mentor, OH) at 14−16 °C. The animal room was equipped with a 24 h light−dark cycle with a light period from 6:00 am to 6:00 pm. Prior to dissection, animals were anesthetized by injection of isotonic 333 mM MgCl_2_ (approximately 50% of body weight) into the body cavity. All reagents were purchased from Sigma–Aldrich (St. Louis, MO) unless otherwise indicated. apATRP was synthesized by ChinaPeptides Co., Ltd.

Electrophysiological techniques were utilized as described previously [24, 25, 27, 33-35, 43, 45]. Briefly, ganglia were desheathed, transferred to a recording chamber containing 1.5 mL of artificial seawater (ASW, 460 mM NaCl, 10 mM KCl, 11 mM CaCl_2_, 55 mM MgCl_2_, and 10 mM HEPES, pH 7.6), continuously perfused at 0.3 mL/min, and maintained at 14−17 °C. Physiological experiments on neuronal excitability were performed in highly divalent (HiDi) saline (368 mM NaCl, 8 mM KCl, 13.8 mM CaCl_2_, 115 mM MgCl_2_, and 10 mM HEPES, pH 7.6), which increases the spiking threshold of neurons and therefore curtails polysynaptic influences. Intracellular recordings were obtained using 5−10 MΩ sharp microelectrodes filled with an electrolyte (0.6 M K_2_SO_4_ plus 60 mM KCl). Grass S88 stimulator was used to provide timing signals for intracellular stimulation. Positive current pulses lasting 3 seconds were used to test excitability of single neurons, with the stimulus interval being 30 seconds. Electrophysiological recordings were digitized online using AxoScope (Molecular Devices, Sunnyvale, CA) and plotted by CorelDraw (Corel Corporation, Ottawa, ON, Canada). Bar graphs were plotted with Prism (version 8, GraphPad Software, La Jolla, CA). Data are expressed as the mean ± S.E.M. All statistical tests were performed using Prism. When the data showed significant effects in ANOVA, individual comparisons were performed with Bonferroni’s correction.

### Microinjection of plasmids

For neurons in the *Aplysia* buccal ganglion that didn’t respond to apATRP, we established two groups. The control neurons were microinjected with the DNA construct pNEX3-EGFP, and the experimental neurons were microinjected with a mixture of pNEX3-EGFP and pNEX3-apATRPR. The EGFP was used as a marker of gene expression. We microinjected the plasmids by pressure injection. The pressure ranged from 20 to 30 psi. To observe the microinjected plasmid volume, we mixed the plasmids with 2% fast green buffer (20 mM HEPES, 200 mM KCl, pH = 7.37) at 1:1. We stopped injecting when the plasmid volume expanded 1/3 of the volume of the injected neuron and the neuron could be seen to turn green due to fast green (see Fig 1B). In the experimental group, if neurons had green fluorescence under an Olympus fluorescence microscope (see Fig 1C), we assumed that these neurons expressed both EGFP and apATRPR.

**Fig 1.**
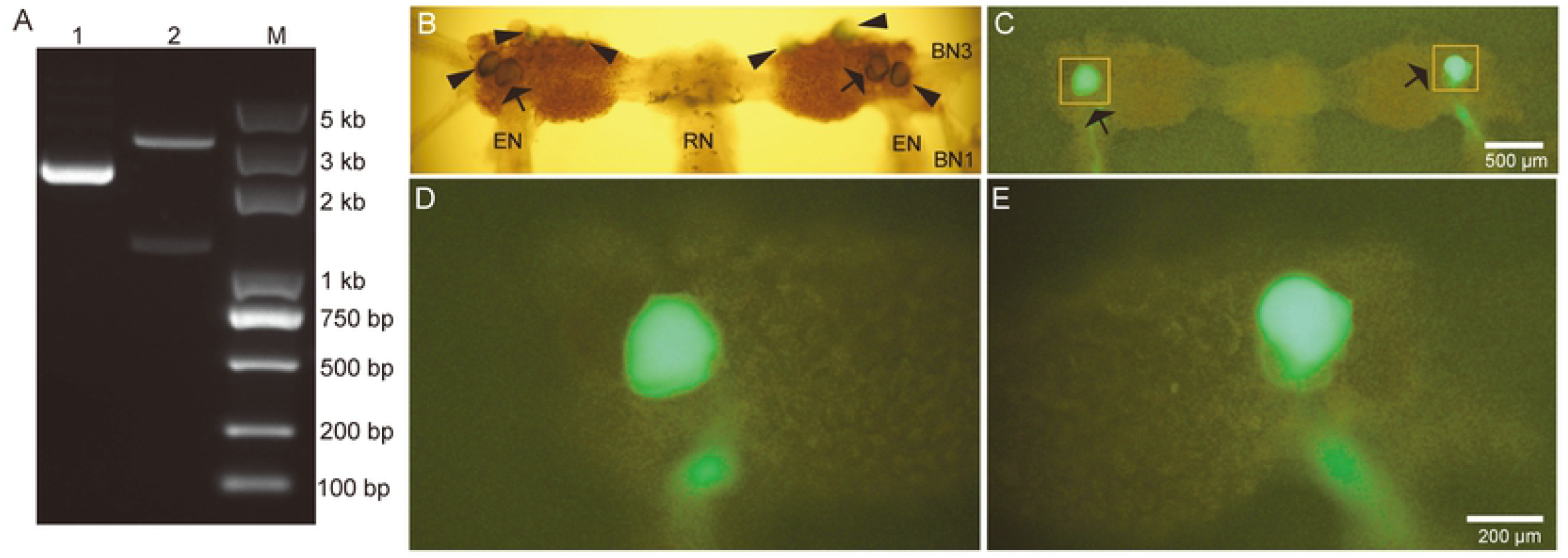
Verification of pNEX3-apATRPR and apATRPR genes, and B1/B2 neurons expressing the receptor apATRPR. (A) Gel electrophoresis of the plasmid pNEX3-apATRPR (approximately 4274 bp) and apATRPR gene (approximately 1215 bp) digested from the BamHI-KpnI restriction fragment of pNEX3 (approximately 3059 bp). apATRPR gene has been verified by sequencing. Lane 1: plasmid pNEX3-apATRPR; lane 2: apATRPR gene (the lower band) and pNEX3 vector (the upper band); M: marker. (B) The caudal surface of a buccal ganglion viewed with a regular light source. B1/B2 and other neurons on the left side were microinjected with plasmids pNEX3-apATRPR and pNEX3-EGFP, and B1/B2 and other neurons on the right side were microinjected with only plasmid pNEX3-EGFP. Arrows and arrowheads: cells with successful injection (fast green). (C) B1/B2 neurons on both sides showed bright fluorescent green (arrows) under a fluorescence microscope. The left B1/B2 neuron expressed apATRPR and EGFP, and the right B1/B2 neuron expressed EGFP. Other neurons marked with arrowheads in (B) did not express injected genes. (D-E) A magnified view of the injected neurons in (C) showing left B1/B2 neuron (D) and right B1/B2 neuron (E). Scale bars in B and C: 500 μm (scale bar in C is for B and C); Scale bars in D and E: 200 μm (scale bar in E is for D and E). EN: esophageal nerve; BN1: buccal nerve 1; BN3: buccal nerve 3; RN: radular nerve.

### Culture of *Aplysia* neurons and detection of plasmid expression

After injection, the buccal ganglia were cultured at 18 °C. The culture medium was made up of an aliquot of *Aplysia* hemolymph and L15 at 1:1 by volume [79, 80]. Then, we added a 1% total volume of 50 mg/ml ampicillin sodium salt (Sigma–Aldrich: A9518-25G-9) solution and a 1% total volume of 200 mM L-glutamine (Sigma–Aldrich: V900419-100G). The culture medium was prepared freshly each time. *Aplysia* hemolymph was prepared from large live *Aplysia* (> 350 gm) using a syringe through a 0.22 μm filter, and aliquots of hemolymph were stored at −80 °C. L-15 medium powder (Leibovitz) (Gibco: 41300039) was supplemented with the salts to make 1-liter solution (L-15 power 13.7 g, NaCl 15.4 g, D-Glucose 6.24 g, MgSO_4_ 3.15 g, KCl 0.35 g, NaHCO_3_ 0.17 g, MgCl_2_.6H_2_O 5.49 g, CaCl_2_ 1.08 g, HEPES 3.53 g, pH, 7.4-7.5; osmolarity: ∼ 1000 mmol/kg). Then, 10 ml of 100 x (5 mg/ml) gentamicin sulfate salt (Sigma–Aldrich: E003632-1G) solution was added. The mixture was filter-sterilized through a 0.22 μm filter, and stored at 4°C.

During ganglia culture, the gene expression was observed using an Olympus fluorescence microscope every day. If the neurons in the control and experimental groups expressed green fluorescence, we then evaluated the activity of apATRP in these neurons using the procedures described in the “Electrophysiology” section.

## Results

### Construction of plasmid pNEX3-ATRPR

The recombinant plasmid, pNEX3-apATRPR, was identified by digestion and the DNA sequencing (Fig 1A). The verified recombinant plasmid was used to microinject *Aplysia* neurons in the experimental group. The plasmid pNEX3-EGFP was microinjected into neurons in the control group.

### apATRP has no effect on neurons B1/B2

To demonstrate that apATRPR might function as an endogenous receptor of apATRP, we sought to find a target neuron that did not natively express apATRPR in the buccal ganglia. We selected a larger neuron, B8 (∼ 150 μm), and examined B8 excitability changes in response to apATRP. apATRP could increase B8 excitability (Fig 2 A-B, F(3, 6) = 14.89, p < 0.01, n = 3), suggesting that B8 might contain a receptor(s) for apATRP. Therefore, B8 neurons were excluded. After testing other neurons, we found that another large neuron B1/B2 (∼ 210 μm) didn’t respond to apATRP (Fig 2 C-D, F(3, 9) = 1.00, p > 0.05, n = 4), and it was used as the target neuron.

**Fig 2.**
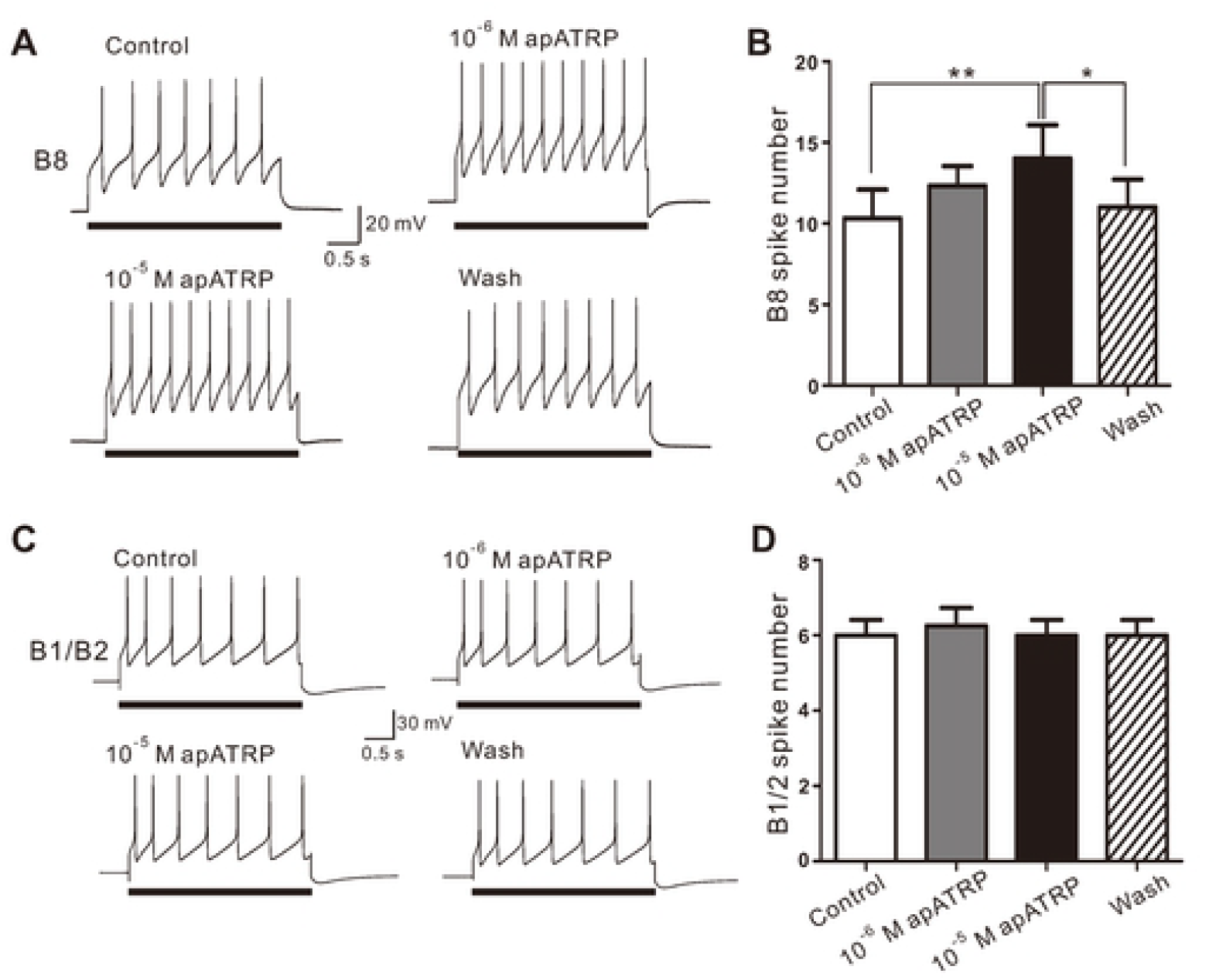
apATRP enhanced B8 excitability but not B1/B2 excitability. (A-B) apATRP increased B8 excitability at 10^−5^ M but not at 10^−6^ M. (A) A representative example. (B) Group data. Bonferroni post hoc tests: * (p < 0.05), ** (p < 0.01). Error bars, SE. (C-D) apATRP had no significant effect on B1/B2 excitability. (C) A representative example. (D) Group data. Bars in A and C denote current injections. Recordings in (A, C) were made in high divalent saline.

### Microinjection of pNEX3-apATRPR into the target neuron

In each hemi-ganglion of the buccal ganglion, there are one B1 and one B2 neuron. Thus, there are four B1/B2 neurons on both sides of the buccal ganglion. We set up two groups: the plasmid pNEX3-EGFP mixture with fast green microinjected into B1/B2 neurons as the control group, and the plasmid pNEX3-EGFP and pNEX3-apATRPR mixture with fast green microinjected into the contralateral B1/B2 neurons as the experimental group. Fast green can be visualized with a regular light source in a microscope, which allowed us to make sure that the plasmid injection was successful (Fig 1B).

After injection, we placed the buccal ganglion into the cell culture for 1-3 days, and observed whether B1/B2 neurons exhibited green fluorescence. If we observed green fluorescence, we considered that neurons were successful in expressing the EGFP in the control group, and co-expressing the apATRPR and EGFP in the experimental group. In 32 cells injected with these plasmids, 30% of the cells subsequently expressed the genes.

### apATRP could enhance the excitability of B1/B2 neurons that exogenously expressed apATRPR

We perfused apATRP into the recording dish and tested B1/B2 excitability. The results showed that B1/B2 excitability in the experimental group was enhanced (Fig 3 A-B, F(3, 9) = 44.84, p < 0.0001, n = 4), and B1/B2 excitability in the control group showed no significant changes (Fig 3 C-D, F(3, 6) = 1.60, p > 0.05, n = 3). The data indicated that the neuropeptide receptor, apATRPR, could mediate the excitability increase in native *Aplysia* neurons in response to its ligand, apATRP.

**Fig 3.**
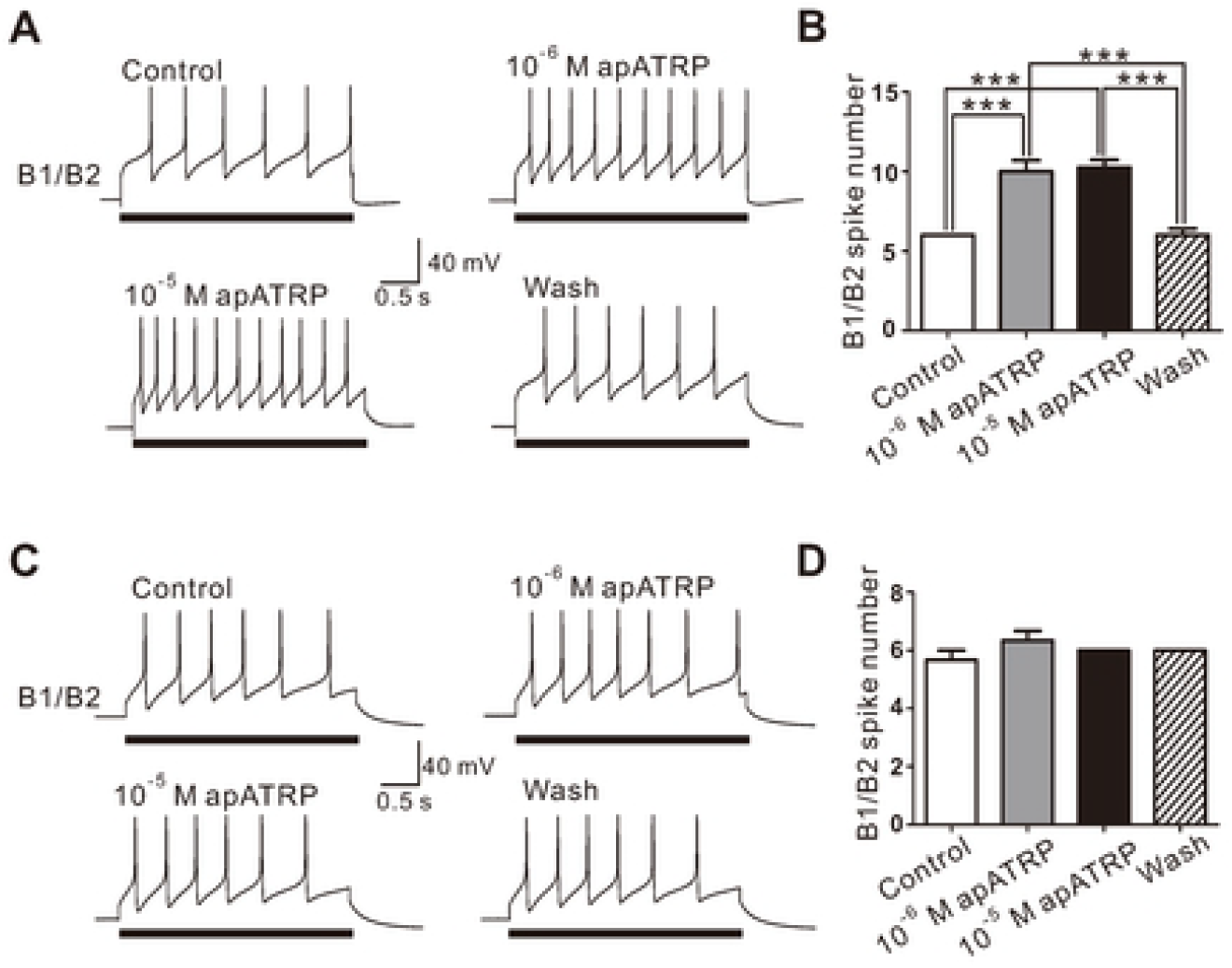
apATRP increased the excitability of B1/B2 neurons that expressed the receptor apATRPR. (A-B) At 10^−6^ M and 10^−5^ M, apATRP increased B1/B2 excitability, which expressed the receptor apATRPR. (A) Representative example. (B) Group data. Bonferroni post hoc tests: *** (p < 0.001). Error bars, SE. (C-D) apATRP had no significant effects on B1/B2 neurons that do not express the receptor apATRPR. (C) A representative example. (D) Group data. Recordings in (A, C) were made in high divalent saline. Bars in A and C denote current injections in B1/B2.

## Discussion

In this work, we have characterized physiological functions of a neuropeptide receptor, apATRPR, expressed in *Aplysia* neurons. Earlier [39, 50], we showed that apATRPR is expressed in the *Aplysia* CNS. Moreover, apATRP and several other ligands can activate apATRPR expressed in a cell line, and these effects matched their enhancing effects on B61/B62excitability. These pieces of evidence are consistent with the notion that apATRPR may mediate the effects of ATRP on B61/B62 neurons. However, prior to the present work, no direct evidence showed that apATRPR could mediate excitability increase in an *Aplysia* neuron. This evidence is critical given two considerations. First, apATRPR activity in a cell line is often measured by a ligand’s ability to increase IP1 concentration in Gαq signaling pathway, whereas excitability increase likely requires an ultimate action of a GPCR on some specific ionic channels [81]. Second, there might be additional ATRP receptor(s) other than the identified apATRPR in *Aplysia* that might mediate excitability increase. Indeed, two allatotropin receptors have been characterized in annelid *Platynereis* [48], which together with mollusc *Aplysia*, belongs to a superphylum: lophotrochozoa.

To provide further evidence, we sought to determine if apATRPR might mediate excitability increase in a neuron that does not express apATRPR. Among several buccal neurons, B1/B2 neurons did not show an excitability increase in response to apATRP. To express exogenous genes in *Aplysia* neurons, we used the plasmid vector pNEX3 [5, 67-69, 75, 76] to construct pNEX3-EGFP and pNEX3-apATRPR, and microinjected B1/B2 neurons in the control group with pNEX3-EGFP, those in the experimental group with both pNEX3-EGFP and pNEX3-apATRPR. B1/B2 neurons in the experimental group could be excited by apATRP, whereas B1/B2 neurons in the control group could not. Thus, our study provides the first evidence that apATRPR indeed mediates excitability increase in a neuron that does not express apATRPR. Taken together with earlier work showing that pNEXδ or pNEX3 can express GPCRs for glutamate or octopamine [64, 69], pNEX, including pNEXδ and pNEX3, proves to be an effective plasmid to express GPCRs for both small molecule transmitters and neuropeptides in *Aplysia* neurons.

We expect that such a procedure could be readily applied to demonstrate physiological functions of neuropeptide receptors in native neurons in model systems with reasonably large identifiable neurons, such as other molluscs, annelids and possibly some arthropods. Notably, compared with invertebrate genetic organisms *C elegans* [82] and *Drosophila* [83], life spans of molluscs and annelids are relatively long, making it difficult to use transgenes to manipulate gene expression. Consequently, the procedure described in this paper should be particularly useful in these animals to study functions of genes in native neurons.

In summary, we provide direct evidence indicating that the neuropeptide receptor, apATRPR, is sufficient to mediate an excitability increase to its ligand, apATRP, in *Aplysia* neurons. This is a further proof that *Aplysia* is an advantageous model system for the study of peptidergic neuromodulation in well-defined neural circuits for feeding, locomotion and other behaviors.

## Acknowledgements

Not applicable.

## Authors’ contributions

**Conceptualization:** Guo Zhang, Bong-Kiun Kaang, Jian Jing

**Data curation:** Guo Zhang, Shi-Qi Guo, Si-Yuan Yin, Wang-Ding Yuan, Ping Chen, Ji-Il Kim, Hui-Ying Wang

**Formal analysis:** Wang-Ding Yuan, Ji-Il Kim, Hui-Ying Wang

**Funding acquisition:** Guo Zhang, Hai-Bo Zhou, Abraham J. Susswein, Bong-Kiun Kaang, Jian Jing

**Methodology:** Shi-Qi Guo, Wang-Ding Yuan, Ping Chen

**Project administration:** Hai-Bo Zhou, Abraham J. Susswein, Bong-Kiun Kaang, Jian Jing

**Supervision:** Abraham J. Susswein, Bong-Kiun Kaang, Jian Jing

**Validation:** Guo Zhang, Shi-Qi Guo, Wang-Ding Yuan, Jian Jing

**Writing-original draft:** Guo Zhang, Abraham J. Susswein, Bong-Kiun Kaang, Jian Jing

